# Identifying Reproducibly Important EEG Markers of Schizophrenia with an Explainable Multi-Model Deep Learning Approach

**DOI:** 10.1101/2024.02.09.579600

**Authors:** Martina Lapera Sancho, Charles A. Ellis, Robyn L. Miller, Vince D. Calhoun

**Affiliations:** Tri-institutional Center for Translational Research in Neuroimaging and Data Science Georgia State University, Georgia Institute of Technology, and Emory University, Atlanta, USA

**Keywords:** Schizophrenia, EEG, Deep learning, Explainable AI, Multi-model analysis

## Abstract

The diagnosis of schizophrenia (SZ) can be challenging due to its diverse symptom presentation. As such, many studies have sought to identify diagnostic biomarkers of SZ using explainable machine learning methods. However, the generalizability of identified biomarkers in many machine learning-based studies is highly questionable given that most studies only analyze explanations from a small number of models. In this study, we present (1) a novel feature interaction-based explainability approach and (2) several new approaches for summarizing multi-model explanations. We implement our approach within the context of electroencephalogram (EEG) spectral power data. We further analyze both training and test set explanations with the goal of extracting generalizable insights from the models. Importantly, our analyses identify effects of SZ upon the α, β, and θ frequency bands, the left hemisphere of the brain, and interhemispheric interactions across a majority of folds. We hope that our analysis will provide helpful insights into SZ and inspire the development of robust approaches for identifying neuropsychiatric disorder biomarkers from explainable machine learning models.

## I. Introduction

Schizophrenia (SZ) is a neuropsychiatric disorder that affects approximately 20 million people globally [1]. However, despite its prevalence, diagnosis and treatment can be difficult because symptoms vary across individuals and overlap with other disorders [2]. As such, data-driven diagnosis based on the identification of brain activity biomarkers, like spectral activity in electroencephalography (EEG) data, could be particularly impactful. Explainable machine learning methods offer an approach for identifying key EEG biomarkers. However, due to the inherent randomness involved in training those models, the features identified as important in a single model may not be generalizable as biomarkers. As such, analyses combining insights from many models are needed. In this study, we train a model for SZ diagnosis on EEG spectral amplitude data. We then present (1) novel approaches for visualizing multi-model explanations and (2) a novel feature interaction explainability approach. Importantly, our approaches identify effects of SZ upon the α, β, and θ frequency bands, the left hemisphere of the brain, and interhemispheric interactions that are reproducibly important in a significant manner across folds.

SZ symptoms can be quite diverse, including positive symptoms like delusions or hallucinations, negative symptoms like a flattening of emotion or loss of pleasure, or general symptoms like loss of cognitive function [1]. These symptoms can vary across individuals and overlap with those of other neuropsychiatric disorders [2]. As such, there has been a growing impetus to identify biomarkers that can enable a more informed empirical diagnosis approach or the development of an automated diagnostic clinical decision support system [3].

Relative to other neuroimaging modalities that have been used in SZ analysis – like functional magnetic resonance imaging (fMRI) [4] or magnetoencephalography (MEG) [2] - EEG is relatively cheap and easy to collect, while also having a high temporal resolution. Many EEG analysis studies use either raw EEG [1], [5], or extracted features [6]. Extracted features offer enhanced explainability [6], which is important if these methods are to be used for biomarker identification. EEG spectral features are commonly used in the analysis of SZ and other neuropsychiatric disorders [6].

Unfortunately, the effects of disorders upon neuroimaging data can be very complex and have led to the use of machine learning and deep learning methods in EEG studies [1], [5], [7]. While these studies are increasingly applying spatial, spectral, temporal, or interaction explainability [7] approaches to explain the patterns uncovered by their models, analyzing a single model or simply showing a distribution of explanations across folds with the hopes of extracting generalizable scientific insights can be problematic given the variability of patterns identified by models resulting from random weight initializations or cross-validation splits. As such, there is a growing need in SZ and broader neuropsychiatric disorder analysis to develop approaches for analyzing multi-model explanations and identifying biomarkers that are reproducibly important. Moreover, most studies only analyze explanations for either training [8] or test data [7], while insights might be had from both. While overfit models learn some noise, training set explanations illuminate a larger portion of the available data and the importance of features for samples in which models obtain near-perfect classification performance. In contrast, test set explanations give insights into subsets of the total available data or patterns that could be more generalizable.

In this study, to identify generalizable SZ biomarkers, we train a multilayer perceptron (MLP) for SZ diagnosis on spectral amplitude information. We then apply explainability approaches to systematically evaluate which features were most important to the model across 25 folds. We identify (1) key frequency bands, (2) key channels, (3) interactions that the model learned between frequency bands, and (4) interactions that the model learned between channels. We further analyze how reproducibly the model relies upon the same features and identifies the same interactions across folds and across training-validation (train-val) and test sets, highlighting those results that are likely to be most reliable and generalizable.

## II. Methods

In this section, we describe our approach. We extract spectral amplitude, train an MLP for SZ diagnosis, obtain spectral and spatial importance, and apply spectral and spatial interaction approaches.

### A. Data Collection

We used a publicly available dataset that was collected at the Institute of Psychiatry and Neurology in Warsaw, Poland [9]. The dataset has been used in multiple previous studies [1], [5]. It has resting state scalp EEG recordings from 14 individuals with SZ (SZs) and 14 healthy controls (HCs) who each gave written informed consent prior to the study. The recordings had a 15-minute duration and 250 Hz sampling rate, and a standard 10-20 electrode format with 64 electrodes.

### B. Data Preprocessing

Like previous studies [6], we used 19 channels: Fp1, Fp2, F7, F3, Fz, F4, F8, T7, C3, Cz, C4, T8, P7, P3, Pz, P4, P8, O1,and O2. To increase the available sample size, we separated the recordings into 25-second epochs using a moving window approach (step size = 2.5 seconds). We next upsampled the HCs to rectify class imbalances by randomly duplicating a subset of samples within each HC. For feature extraction, we fast Fourier transformed (FFT) the data, calculated the amplitude at each frequency, and averaged within each frequency band. Frequency bands included: δ (0 – 4 Hz), θ (4 - 8 Hz), α (8 – 12 Hz), β (12 – 25 Hz), γlower (25 – 55 Hz), and γupper (65 – 125 Hz). We obtained 114 features (6 frequency bands x 19 channels) and feature-wise z-scored them.

### C. Model Development

We developed an MLP with 3 dense layers (36 nodes, 18 nodes, 2 nodes) for SZ versus HC classification. The first two layers had rectified ReLU activations and 30% dropout. The final layer had a softmax activation. We trained the MLP with a nested group shuffle split cross-validation (CV) approach to ensure that samples from a subject were not mixed across the training, validation, and test sets. We used 25 outer folds (80%= train-val, 20% test) and 10 inner folds (80% training, 20% validation). We used class-weighted binary cross-entropy loss and an Adam optimizer with a learning rate optimized in the inner folds via a grid search over [1e-05, 5e-05, 1e-04, 5e-04, 0.001, 0.005, 0.01, 0.05, 0.1, 0.5]. We trained the models for 200 epochs with a batch size of 128 and shuffling after each epoch. We used checkpoints to select the inner-fold model from the epoch with the maximal F-measure (F) - geometric mean between validation sensitivity (SENS) and specificity (SPEC). We measured performance with F, SENS, SPEC, accuracy (ACC), balanced ACC (BACC), F1 score (F1) and precision (PREC). We then calculated their mean (μ) and standard deviation (σ) across folds.

### D. Spectral and Spatial Approach

For explainability, we computed Layer-Wise Relevance Propagation (LRP) with the iNNvestigate toolbox [10]. LRP outputs feature relevance by assigning a total relevance value of 1 to the output node of a target class and backpropagating that relevance through the model using a relevance rule [7]. LRP relevance can be positive or negative (i.e., when a feature supports the assignment of a sample to the target class or an off-target class) [7]. In this study, we used the αβ relevance rule (α=1, β=0) to only output positive relevance [7]. After outputting relevance for each train-val and test sample, we normalized the absolute relevance for each sample making their total absolute relevance sum to 1. We then obtained spectral and spatial relevance by summing the relevance for features in each frequency band and each channel, respectively. We performed one-sample t-tests in each train-val and test set comparing the relevance distribution for each feature across samples to the μ relevance across feature groups (i.e., 1/19 for channels and 1/6 for frequency bands). We applied False Discovery Rate (FDR, α=0.05) correction to account for multiple comparisons for each fold. We then calculated the μ relevance for each feature across folds and the percentage of folds with significant results.

### E. Spectral and Spatial Interaction Approach

We next explored feature interactions with a novel approach based on [7], performing separate analyses for frequency band interactions and channel interactions. We (1) calculated the relevance for each feature group, as in the previous section, (2) perturbed a group of features by replacing them with zeros, (3) calculated the total absolute relevance for each group for the perturbed samples, and (4) compared the relevance for each group before versus after perturbation. For insight into the consistency across models and reproducibility across datasets of the interactions, we used unpaired two-sample t-tests comparing relevance before and after perturbation followed by FDR (α=0.05) correction on both train-val and test datasets within each fold. We then calculated the percentage of significant folds for each feature. This analysis gave insight into inter-spatial and inter-spectral interactions. For example, if F7 relevance decreased after the perturbation of O2, then the information in O2 likely enabled the predictive power of F7.

## III. Results and Discussion

In this section, we describe and discuss our findings along with the limitations and future directions of our methodology.

### A. Model Performance

Table I shows the µ and σ of our test performance across folds. Performance is above 70% across metrics, and SENS and SPEC are comparable. Our performance exceeds that of other studies [6], and while some studies have higher performance, many do not employ a subject-wise CV approach [5]. Our training performance is near 100% in all metrics across folds.

**TABLE 1.** Model Performance Metrics.

| Across models | Test Performance Metrics |  |  |  |  |  |  |
| --- | --- | --- | --- | --- | --- | --- | --- |
|  | ACC | BACC | SENS | SPEC | F1 | F | PREC |
| $\mu$ | 0.73 | 0.75 | 0.77 | 0.74 | 0.70 | 0.71 | 0.68 |
| $\sigma$ | 0.09 | 0.09 | 0.18 | 0.16 | 0.13 | 0.05 | 0.20 |

### B. Spectral and Spatial Importance Results

Fig. 1A and 1B show the spectral importance for the trainval and test data, respectively. In both analyses, α, β, and θ are most important and significant in more than 50% of models. In both cases, α has the highest μ relevance and is significant in all folds. The α-band is closely linked to clinical SZ symptoms [11]. The next most important band differs between analyses, with θ and β having second-highest μ relevance in train-val and test analyses, respectively. While the portion of folds with β significance increased from 72% in the train-val set to 80% in the test-set, for θ significance, values decreased from 84% to 68%. Importantly, SZ is linked to aberrant cholinergic mechanisms that commonly induce abnormal β and θ oscillations [12].

**Fig. 1.**
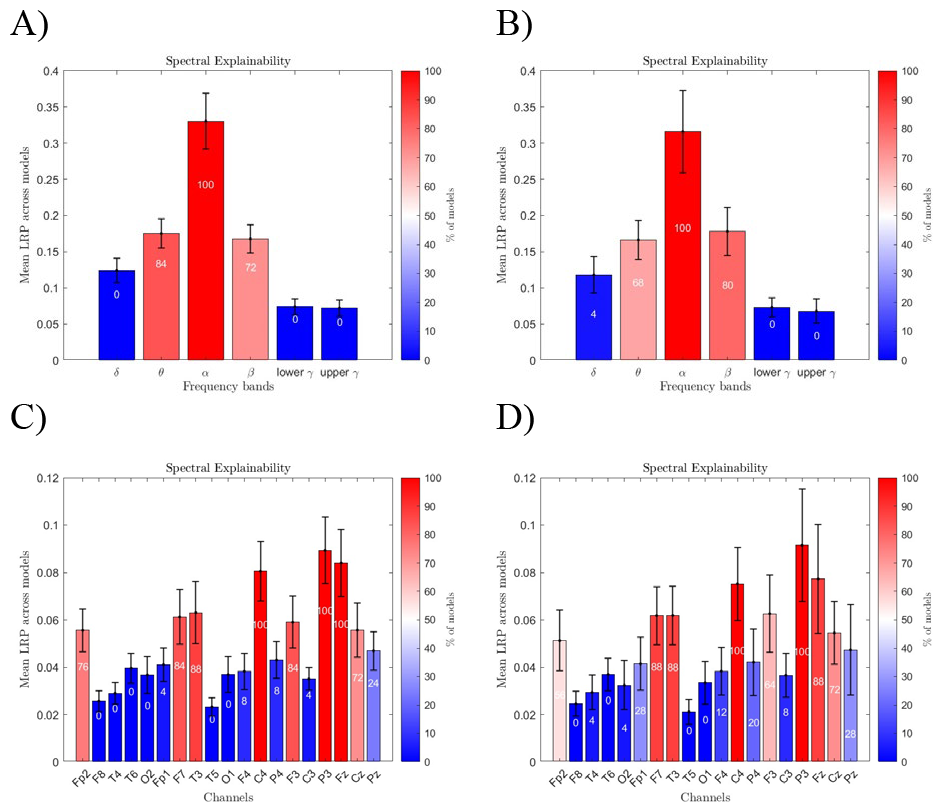
Spectral and spatial importance results. A) and B) show spectral importance results on train-val and test data, respectively. C) and D) show spatial importance results on train-val and test data, respectively. Bar heights depict μ and σ relevance for each feature across folds. The color scale represents the percentage of folds in which a feature has above-average relevance (specific percentages are included inside each bar).

Fig. 1C and 1D show the spatial importance for the trainval and test sets, respectively. Channels T3, C4, F3, P3, Fz, Fp2, F7 and Cz are significant in more than 50% of models in both analyses. In both cases, P3 has the highest µ relevance followed by Fz, and C4. Interestingly, P3 and C4 have significant relevance in all folds for the train-val analysis and most folds for the test analysis and are associated with the frontoparietal network that is disrupted in SZs due to intranetwork dysconnectivity [13]. The next μ relevance rankings among channels then differ between analyses. For train-val, C4 is followed by T3, F7, F3, Cz, and Fp2. However, for testing, C4 is followed by F3, T3, F7, Cz, and Fp2. Interestingly, F3, Fz and Fp2 are significant in fewer test sets than train-val sets. Cz and T3 significance across folds remained constant from train-val to test analyses, while that of F7 increased. Interestingly, T3, F7, F3 and P3 are all located within the left hemisphere, which has been related to negative SZ symptom severity [14]. We can therefore conclude that the spatial features most generalizably relevant to SZs are P3, Fz, C4, T3, F7, F3, Fp2, and Cz.

Importantly, although percentages within individual feature groups vary from train-val to test analyses, no feature with significance in more than 50% of folds decreased to having significance in less than 50% of folds between analyses. This indicates that, although our models slightly overfit the train-val data (ACC decreased around 27%), both spectral and spatial explanations were highly generalizable across datasets.

### C. Spectral and Spatial Interaction Results

Fig. 2A and 2B show the spectral interaction results for train-val and test sets, respectively. Unfortunately, the percentage of folds with frequency bands with decreased relevance is relatively small across analyses, although a few folds identify δ-θ and δ-α interactions. This may be because the 19 channels associated with each frequency band had varying effects when perturbed or responding to perturbation.

**Fig. 2.**
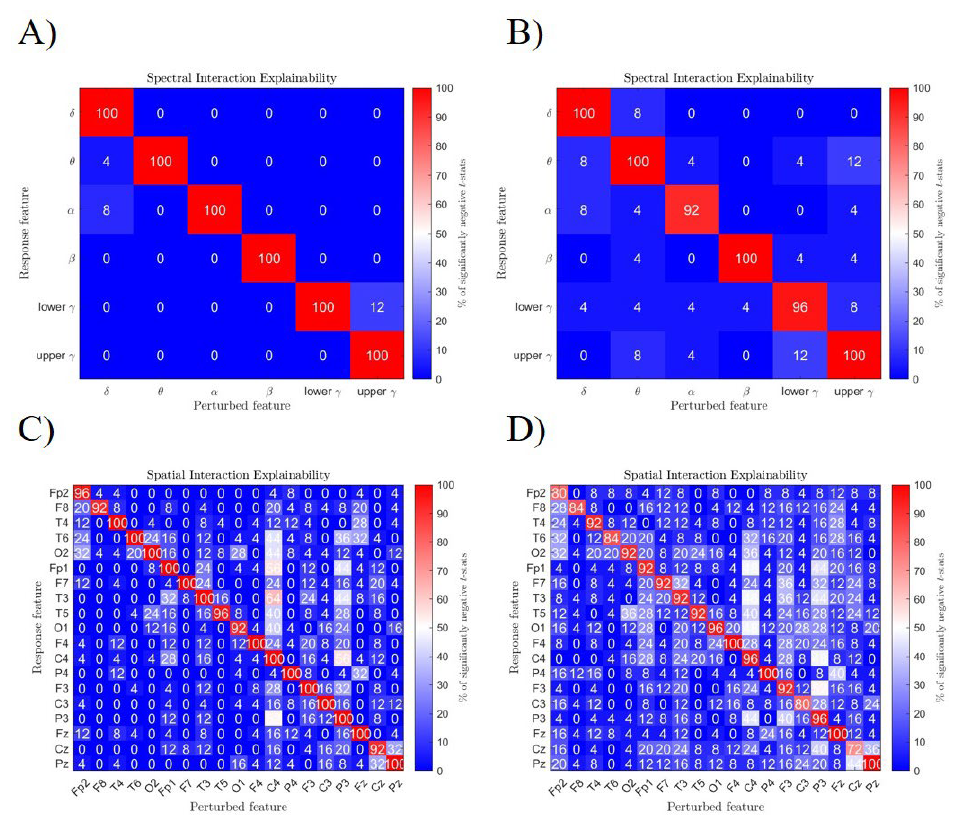
Spectral and spatial interaction results. A) and B) show spectral interaction results on train-val and test data, respectively. C) and D) show spatial interaction results on train-val and test data, respectively. The color scale shows the percent of folds with significantly decreased relevance (negative t-statistic, p<0.05) after our perturbation approach.

Fig. 2A and 2B show the spatial interaction results for the train-val and test data, respectively. We observe strong interhemispheric interactions, which fit with existing studies [15]. Namely, C4 (right hemisphere) significantly interacts with F7, P3, and Fp1 (left hemisphere) in more than 50% of the train-val sets and nearly 50% of the test sets. P3 has significant C4 interaction in more than 50% of the train-val sets. Interestingly, P3 also interacts with F3 in 52% of folds and is the only interaction found in more than 50% of the test sets.

This could be related to disrupted working memory in SZs [1] that is correlated with frontoparietal θ and α coherence [16]. Fp1, T3 and F3 are also highly interactive, which fits with findings that have linked reduced fronto-temporal connectivity to internal voice symptoms in SZ [17]. Generally, the test sets had more interactions, but the train-val sets were more consistent across folds (i.e., more concentrated spatially and present in more folds).

### D. Limitations and Future Work

Although our model performance is above chance-level and serves as a solid basis for our explainability analysis, our model architecture and hyperparameter tuning strategies could be improved to yield higher model performance. In our interaction approach, we replaced features with zeros. This perturbation risks creating out-of-distribution samples that may yield invalid explanations. An alternative, though more computationally expensive, would be the implementation of a permutation-based perturbation approach with a sufficiently high number of iterations. Increasing the number of training folds or training the model with multiple initializations could also improve insights in future studies.

## IV. Conclusion

Many studies have sought to identify diagnostic biomarkers of SZ due to its diverse symptom presentation. However, the findings of machine learning studies can be of questionable generalizability, as they can be black-box or yield explanations that consider relatively few folds. Analyzing explanations across many machine learning models could lead to the identification of more generalizable biomarkers. In this study, we present (1) a novel feature interaction-based explainability approach and (2) several new approaches for summarizing multi-model explanations. We demonstrate the applicability of our approaches within the context of EEG-based SZ diagnosis with extracted spectral features. Our models obtain a high level of performance and identify effects of SZ on the α, β, and θ frequency bands, the left hemisphere of the brain, and interhemispheric interactions. We hope that our analysis will provide helpful insights into SZ and inspire the development of robust approaches for identifying neuropsychiatric disorder biomarkers from explainable machine learning models.

## Notes

https://journals.plos.org/plosone/article?id=10.1371/journal.pone.0188629

### Competing Interest Statement

The authors have declared no competing interest.

